# Correlative single-molecule fluorescence barcoding of gene regulation in *Saccharomyces cerevisiae*

**DOI:** 10.1101/2020.08.12.247643

**Authors:** Sviatlana Shashkova, Thomas Nyström, Mark C Leake, Adam JM Wollman

## Abstract

Most cells adapt to their environment by switching combinations of genes on and off through a complex interplay of transcription factor proteins (TFs). The mechanisms by which TFs respond to signals, move into the nucleus and find specific binding sites in target genes is still largely unknown. Single-molecule fluorescence microscopes, which can image single TFs in live cells, have begun to elucidate the problem. Here, we show that different environmental signals, in this case carbon sources, yield a unique single-molecule fluorescence pattern of foci of a key metabolic regulating transcription factor, Mig1, in the nucleus of the budding yeast, *Saccharomyces cerevisiae*. This pattern serves as a ‘barcode’ of the gene regulatory state of the cells which can be correlated with cell growth characteristics and other biological function.

**Highlights:**

- Single-molecule microscopy of transcription factors in live yeast
- Barcoding single-molecule nuclear fluorescence
- Correlation with cell growth characteristics
- Growth in different carbon sources

## Introduction

Cell survival depends on adaptation to environment. Cells sense and respond to extracellular signals through complex signalling pathways [1,2]. In most processes, signal response is achieved by adjusting protein levels through degradation, RNA processing or through families of protein transcription factors (TFs) [3] which enter the nucleus, find target sites within genes and either repress or enhance gene expression. Each TF binding sequence is short, typically a few 10s of base pairs, and so non-specific that the sequence will typically occur over thousands of times in the genome, more than nearly any factor can occupy [4]. The key problem of predicting where specific TFs will sit in the genome and which genes are regulated has been termed the ‘futility paradox’ [5]. Recent new techniques may provide the solution, chromosome conformation capture techniques have started to reveal the 3D arrangement of genes in the nucleus[6] while single-molecule microscopy can now follow single TFs as they bind to DNA [7].

Single-molecule fluorescence microscopy has been used to image TFs directly in mammalian cells [8,9], showing that TFs speed up searching for their binding sites by sliding along and hopping between DNA filaments. However these commonly used single-molecule imaging methods, such as PALM, exploit very sparse labelling to resolve individual molecules, with fewer than 1% of TFs fluorescently tagged [9], and thus cannot capture the position of all TFs or their molecular architecture. We showed previously, that we could use Slimfield microscopy combined with full protein labelling to capture information about the position of all TFs in a cell nucleus [10]. By using budding yeast cells expressing an endogenously GFP-tagged TF in the glucose repression pathway, Mig1, we showed that this TF and others operate as clusters and that we could observe the ultra-structure of these clusters in the nucleus. This ultra-structure resulted in a pattern of nuclear foci with a distribution of molecular stoichiometries.

In the budding yeast *Saccharomyces cerevisiae*, genes required for metabolism of alternative carbon sources are controlled via the glucose repression pathway [11]. In a glucose-rich environment, the transcriptional repressor Mig1 together with the global co-repressor complex occupies target promoters and represses the expression of genes essential for utilisation of sucrose (*SUC2*) [12], galactose (*GAL*) [13] and maltose (*MAL*) [14]. In response to glucose limitation Mig1 no longer associates with the corepressor complex and the repression of the target genes is released and Mig1 translocates out of the nucleus.

Our previous study revealed a pattern of Mig1 clusters in the nucleus in glucose rich conditions, resulting in a distinct distribution of foci stoichiometry [10]. We hypothesised that this pattern directly relates to the specific repressed genes in response to extracellular signals such as different carbon sources. Little is known about Mig1’s behaviour under different carbon sources and how this is correlated to cellular growth characteristics, such as growth rates, doubling times and yield. Here, we performed Slimfield imaging of Mig1-mGFP in yeast cells exposed to different carbon sources. We obtained a specific single-molecule stoichiometry distribution for each input carbon source, confirming our hypothesis and allowing us to ‘barcode’ the gene regulatory state and correlate it with biological function through measured growth rates.

## Materials and methods

### Growth conditions and media

Cells from frozen stocks were pre-grown on standard YPD medium plate (20 g/L Bacto Peptone, 10 g/L Yeast Extract) supplemented with 4% glucose (w/v) at 30°C. For liquid cultures, cells were grown in Yeast Nitrogen Base (YNB) medium (1x Difco™ YNB base, 1x Formedium™ complete amino acid Supplement Mixture, 5.0 g/L ammonium sulphate, pH 5.8-6.0) supplemented with a carbon source of a desired concentration, at 30°C, 180 rpm.

### Growth rates experiments

We have previously shown, that the presence of Mig1-mGFP and Nrd1-mCherry endogenously expressed fluorescent fusions does not affect the growth rate of *S. cerevisiae* [10]. Therefore, for the growth experiments, we used the BY4741 wild type cells pre-grown overnight in YNB complete medium supplemented with 4% glucose (w/v), 30°C, 180 rpm. Cells were then washed to remove any remaining glucose and diluted to OD_600_ ∼0.01 in YNB complete supplemented with one of the following: 4% glucose, 0.2% glucose, 2% galactose, 2% sucrose or 3% ethanol. Obtained cultures were placed onto a 24-well plate, 1.6 ml/well. Media with appropriate carbon sources but without any cells were used as a negative control. Absorbance at 600 nm was measured for more than 60 hours every 45 min under typical growth conditions,30°C, constant shaking (BioTek Synergy™ 2 Microplate Reader, ThermoFisher Scientific). To calculate the growth parameters, the data was analysed with a desktop growth data processing tool, PRECOG [15].

### Slimfield Microscopy

For microscopy experiments, BY4741 cells expressing Mig1-mGFP and Nrd1-mCherry protein fusions were pre-grown overnight in YNB media with 40 g/L glucose, sub-cultured and grown until mid-logarithmic phase, *OD*_*600*_ *∼0*.*7*. Cells were then washed to remove glucose, placed into media supplemented with 4% glucose, 2% sucrose (w/v), 2% galactose, 3% ethanol or 0.2% glucose for 2 h at 30°C, 180 rpm.

For imaging, cells were immobilized by placing 5 µL of the cell culture onto a 1% agarose pad perfused with YNB supplemented with an appropriate carbon source. The pad with cells was sealed with a plasma-cleaned BK7 glass microscope coverslip (22×50 mm).

Cells were imaged with a bespoke Slimfield microscope used previously [10]. Slimfield excitation was implemented via 50mW 488 nm and 50mW (attenuated to 5mW) 561nm wavelength laser (Coherent Inc., OBIS lasers, SantaClara, California, USA), for GFP and mCherry imaging respectively, de-expanded to direct a 10μ m at full width half maximum beam onto the sample at 20mW excitation intensity to observe single GFP in living yeast cells [10]. Fluorescence emission was captured by a 1.49 NA oil immersion objective lens (Nikon). Images were collected at 5 ms exposure time by the EMCCD camera (iXon DV860-BI, Andor Technology, UK) using 80 nm/pixel magnification.

The focal plane was set to mid-cell height using the brightfield appearance of cells. As photobleaching of mGFP proceeded during Slimfield excitation distinct fluorescent foci could be observed with half width at half maximum 250-300 nm, consistent with the diffraction-limited point spread function of our microscope system, which were tracked and characterized in terms of their stoichiometry and apparent microscopic diffusion coefficient. Distinct fluorescent foci that were detected within the microscope’s depth of field could be tracked for up to several hundred ms, to a super-resolved lateral precision ∼40 nm [16] using a bespoke single particle tracking software written in MATLAB (MATHWORKS) and adapted from previously described live cell single-molecule studies [10,17].

The molecular stoichiometry for each track was determined by dividing the summed pixel intensity values associated with the initial unbleached brightness of each foci by the brightness corresponding to that calculated for a single fluorescent protein molecule measured using a step-wise photobleaching technique [18]. Maturation effects of fluorescent protein fusions withing living cells were characterized on similar yeast cell lines previously, indicating typically less than 10-15% immature ‘dark’ fluorescent protein [19].

## Results and discussion

We performed single-molecule Slimfield microscopy on yeast cells expressing Mig1-mGFP and Nrd1-mCherry as a nuclear marker (Fig. 1B) in different carbon sources. In 4% glucose, we observed Mig1 largely present in the nucleus as before [10], as well as the presence of distinct foci within the nucleus corresponding to clusters and assemblies of clusters of Mig1 within the diffraction limit of our microscope. We segmented for the cell and nuclei using threshold based segmentation of the GFP and mCherry images respectively, with a threshold defined as the upper full width at half maximum of the background intensity. Slimfield movies were run through our automated foci detection and tracking software [16], to characterise the intensity of nuclear foci as a function of time. We performed stoichiometry analysis of these intensity traces (Fig. 1C) using a simplified method from our previous works [10,20]. Intensity traces initially resemble exponential decay, as many fluorophores stochastically photobleach. Towards the end of the photobleach, individual fluorophore photobleach events become detectable, with the inset in Fig. 1C showing the final photobleach step before the foci is completely bleached. The intensity of these final steps was pooled and the distribution is shown in Fig. 1D. We quantified the characteristic photobleach intensity of these steps as the peak of the distribution in Fig. 1D to be 3700±600 detector counts. We could then quantify the stoichiometry of the foci i.e. the number of Mig1-mGFP molecules present within the cluster or assembly, by dividing the initial intensity of the foci (as determined by a linear fit to the beginning of the intensity trace) by the characteristic intensity.

**Figure 1:**
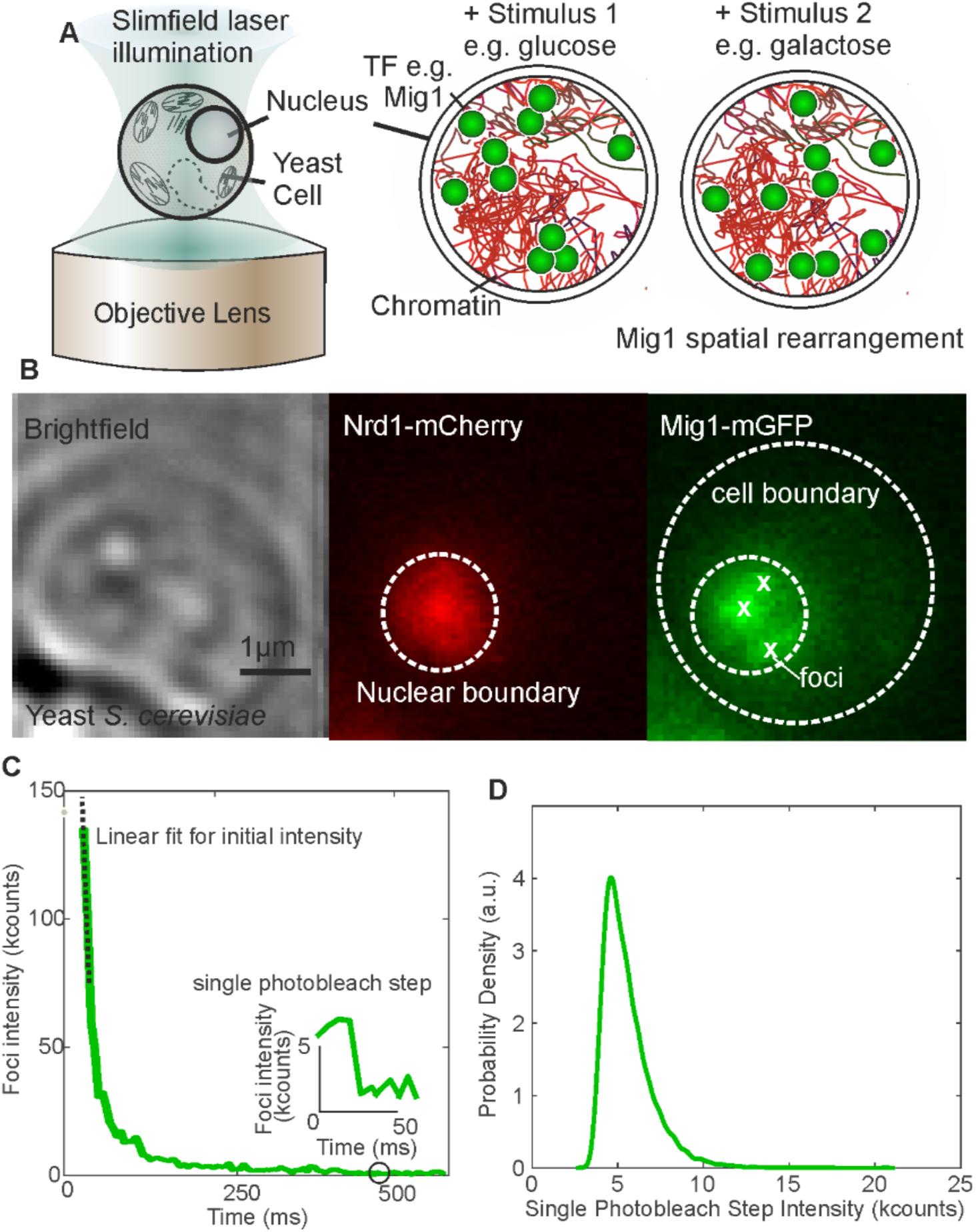
A. Schematic of Slimfield microscopy of yeast cell and the molecule re-arrangement of Mig1 in the nucleus in response to different external stimuli. B. Example brightfield micrograph of a yeast cell and associated Slimfield fluorescence micrographs of nuclear marker Nrd1-mCherry (red) and Mig1-mGFP (green) with cell and nuclear boundaries shown in white dotted lines and detected foci shown as white crosses. C. An example intensity vs time trace for a foci detected in B. Foci rapidly photobleaches with the inset showing the intensity just before bleaching as a distinct step. The initial linear section of the photobleach trace is fitted with a line to determine the initial intensity. D. The distribution of intensity towards the end of all photobleach traces, the peak of which gives the characteristic intensity of a single fluorophore.

We hypothesised that the intensity distribution of foci within the nucleus would provide a unique molecular barcode for the gene regulatory state of the cell population. In other words, we hypothesised that in response to different related external stimuli, the stoichiometry distribution of Mig1 within the nucleus would change, reflecting the different pattern of gene regulation and the re-arrangement of Mig1 onto different sets of target genes. We tested this idea by performing Slimfield microscopy of Mig1-mGFP expressing yeast cells exposed to a range of different carbon sources: high glucose, low glucose, ethanol, galactose and sucrose (Fig. 2). We found statistical differences in the mean stoichiometry in the nucleus between conditions (Fig. 2A) with completely different distributions of stoichiometries (Fig. 2B-F).

**Figure 2:**
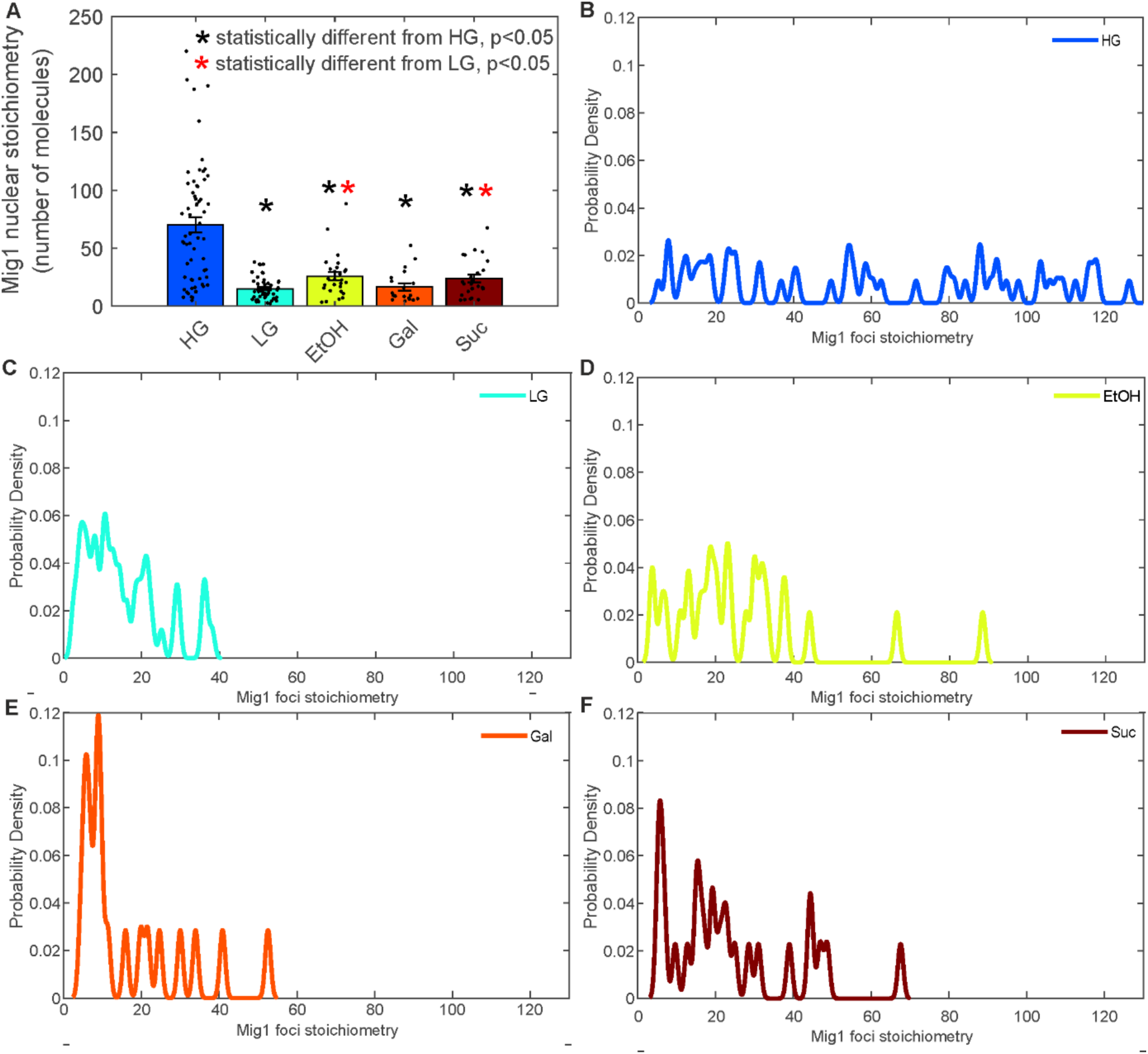
A. Jitter plot of the stoichiometry of foci detected in each of the carbon source conditions (HG – high glucose, 4% w/v; LG – low glucose, 0.2% w/v; EtOH – 3% ethanol; Gal – galactose, 2% w/v; Suc – sucrose, 2% w/v). Mean ± standard error of the mean shown as bars. * indicate statistical differences by students t-test at p<0.05 level from either the high or low glucose condition (black and red respectively). B-F. Distribution of detected foci from A rendered as kernel density estimates.

We sought to determine the biological significance of these stoichiometry distributions or molecular barcodes, by correlating them with biological function, specifically growth characteristics in each condition (Fig. 3). We monitored cultures for more than 60 h and measured the growth curves based on the absorbance at a wavelength of 600 nm (Fig. 3A). When glucose is available in the media, yeast cells undergo fermentative lifestyle despite oxygen presence, and processes including gluconeogenesis, respiration and utilisation of alternative carbon sources are suppressed. The Mig1 protein, as well as a number of other transcription factors of the glucose repression pathway, regulate genes essential for metabolism of non-glucose energy sources. Thus, higher mean apparent stoichiometry of Mig1 in the nucleus (Fig. 2B) in HG likely corresponds to higher target promoter occupancy and repression and is consistent with our previous observations [10]. As glucose is one of the most preferred energy sources for *S. cerevisiae*, cells grown in glucose media have a relatively short lag phase (Fig. 3B) and a higher rate of biomass accumulation compared to non-glucose conditions (Fig. 3C).

**Figure 3:**
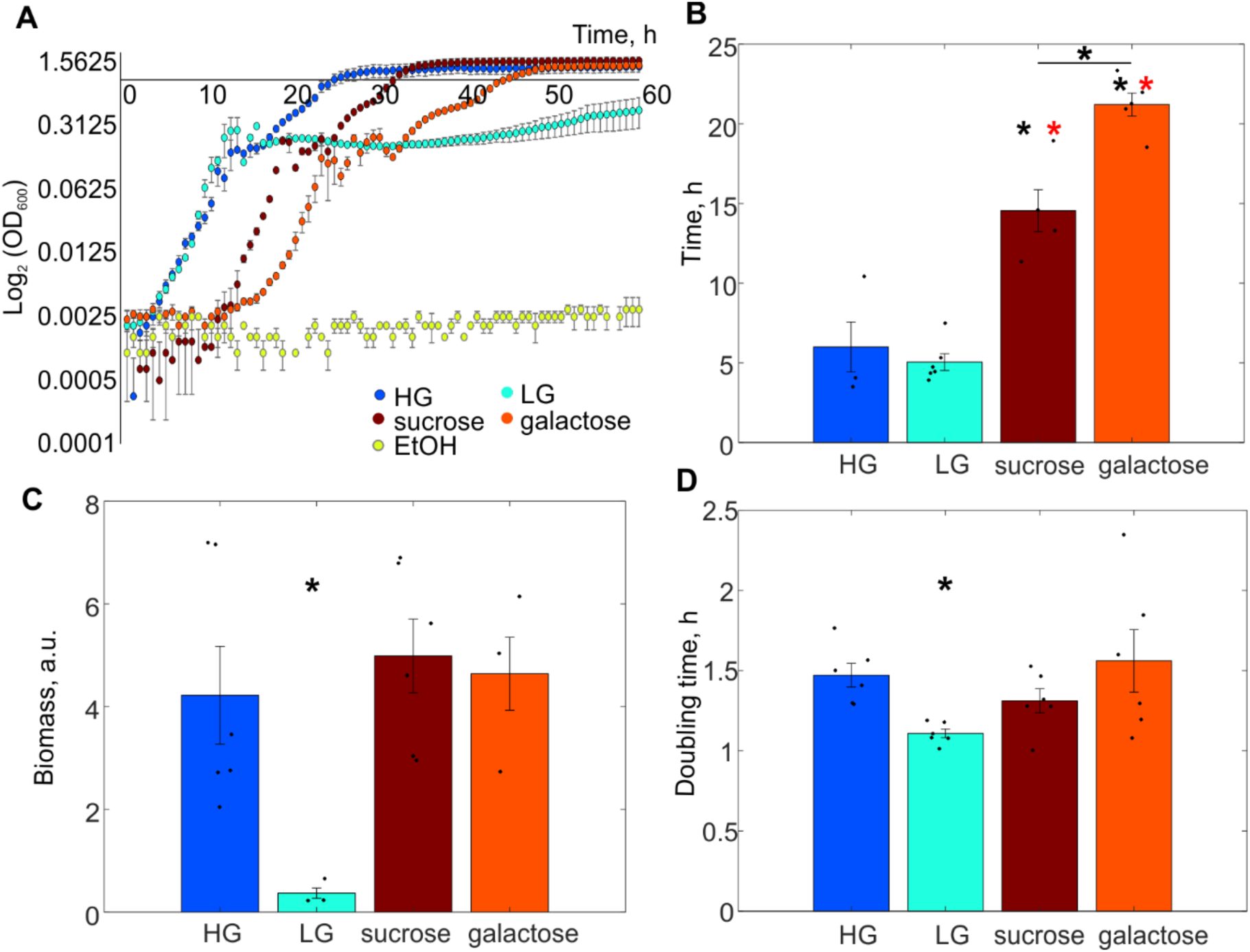
A. Growth curves of yeast cells pre-grown in glucose and sub-cultured into fresh media with various carbon source conditions (HG – 4% glucose w/v, LG – 0.2% glucose w/v, 2% sucrose, 2% galactose or 3% ethanol). Error bars represent standard error. B. Average time of the lag phase in different conditions. Error bars represent standard error. Student’s t-test, p<0.05 from HG or LG (black or red, respectively). C. Average biomass production, yield, obtained by the growth in different conditions Error bars represent standard error of mean. Student’s t-test, p<0.05. D. Mean generation time of cells grown in different carbon source conditions. Standard error of mean, Student’s t-test, p<0.05.

Low concentration of glucose in the environment does not appear to affect the time cells need before they enter the stationary phase (Fig. 3A, B), but results in low yield (Fig. 3C). This can be explained by rapid consumption of all glucose available. Upon glucose limitation, Mig1-induced gene repression is largely released, and the protein mainly relocalises to the cytoplasm [10,17] although some foci remain (Fig. 2C). This corresponds to significantly lower mean stoichiometry (Fig. 2A) as well as more narrow stoichiometry distribution of the nuclear Mig1 compared to that in HG (Fig. 2C).

We were unable to detect any growth on 3% ethanol (Fig. 3A), the only nonfermentable substrate in our study. Indeed, multiple studies that involve ethanol-grown yeast cultures in minimal media, used either a YPE medium (a standard YPD with ethanol instead of glucose) or a glycerol-ethanol mixture [21–24]. However, we did not change the conditions to be consistent with our single-molecule experiments. Therefore, no lag phase time, yield or generation time was estimated for cells in ethanol. Cells grown in sucrose and galactose need significantly more time before the culture enters the exponential growth phase (Fig. 3A, B). However, the yield is comparable to that from cells grown in HG (Fig. 3C). No significant differences in doubling times were detected except for LG (Fig. 3D). This is surprising, and probably comes from our experimental setup limitations. Normally, we would expect the proliferation rate in LG to be similar to HG [10,25].

Despite the differences in Mig1 stoichiometry distributions and means between all conditions (Fig. 2), there is a clear pattern for HG which makes it a distinct feature of glucose repression – there are significantly more and larger Mig1-mGFP foci. Moreover, we observe similar stoichiometry distributions in cells grown in galactose and sucrose (Fig. 2E,F) which corresponds to long lag phases (Fig. 3B). Hence, the time cells need to adapt to the surrounding before they start dividing, seems to be one of the characteristics that could be linked to expression of genes affected by present conditions.

Here, we have demonstrated that single-molecule fluorescence microscopy can be used to ‘barcode’ the state of gene regulation of a specific TF, Mig1. These barcoded states correlate to distinct biological outcomes, characterised here by the growth rates and lag times. The molecular barcoding technique presented here, could be further correlated with other biological readouts, such as the mRNA levels of the specific regulated genes. This could be measured using qPCR and correlated with the single-molecule barcode, allowing transcription profiles to be identified using microscopy of individual cells without having to extract RNA. New microscopy techniques are increasing the resolution possible and moving into 3D [7,26–28], thus the discovery here of unique spatial patterns or barcodes of a TF within the nucleus in response to different stimuli, paves the way for the eventual use microscopy to determine regulated genes.

## Abbreviations

EMCCD: electron multiplying charge-coupled device
GFP: green fluorescent protein
HG: high glucose
LG: low glucose
PALM: photoactivated localisation microscopy
TF: transcription factor
YNB: yeast nitrogen base
YPD: yeast extract peptone dextrose.

## Acknowledgements

We thank Karl Persson, Department of Chemistry and Molecular Biology, University of Gothenburg, for his assistance with the PRECOG software.

## Supplementary

